# Activin signaling informs the graded pattern of terminal mitosis and hair cell differentiation in the mammalian cochlea

**DOI:** 10.1101/380055

**Authors:** Meenakshi Prajapati-DiNubila, Ana Benito-Gonzalez, Erin J. Golden, Shuran Zhang, Angelika Doetzlhofer

## Abstract

The mammalian auditory sensory epithelium has one of the most stereotyped cellular patterns known in vertebrates. Mechano-sensory hair cells are arranged in precise rows, with one row of inner and three rows of outer hair cells spanning the length of the spiral-shaped sensory epithelium. Aiding such precise cellular patterning, differentiation of the auditory sensory epithelium is precisely timed and follows a steep longitudinal gradient. The molecular signals that promote auditory sensory differentiation and instruct its graded pattern are largely unknown. Here, we identify Activin A as an activator of hair cell differentiation and show, using mouse genetic approaches, that a local gradient of Activin A signaling within the auditory sensory epithelium times the longitudinal gradient of hair cell differentiation. Furthermore, we provide evidence that Activin-type signaling regulates a radial gradient of terminal mitosis within the auditory sensory epithelium, which constitutes a novel mechanism for limiting the number of inner hair cells being produced.

## INTRODUCTION

Housed in the inner ear cochlea, the auditory sensory organ contains a spiral shaped sensory epithelium specialized to detect and transduce sound. Along its longitudinal axis, two types of mechano-receptor cells, so called inner and outer hair cells, are arranged in distinct rows, with three rows of outer hair cells and a single row of inner hair cells. To ensure the highly stereotyped arrangement of hair cells, cell cycle withdrawal and differentiation within the auditory sensory epithelium occurs in a spatially and temporally highly coordinated manner. Auditory sensory progenitors (pro-sensory cells) exit the cell cycle in an apical-to-basal gradient (Ruben, 1967; Chen and Segil, 1999; Lee et al., 2006), whereas their differentiation into hair cells and supporting cells occurs in an opposing, basal-to-apical gradient (Chen et al., 2002). Previous studies revealed that the morphogen Sonic Hedgehog (SHH) plays a key role in setting up the spatial and temporal pattern of auditory hair cell differentiation. In the undifferentiated cochlea, pro-sensory cells are exposed to high levels of SHH secreted by the adjacent spiral ganglion neurons (SGNs) (Liu et al., 2010; Bok et al., 2013). Ablation of SHH in SGNs or cochlear epithelial-specific ablation of co-receptor Smoothened results in premature hair cell differentiation and in the most extreme case a reversal of the gradient of hair cell differentiation (Bok et al., 2013; Tateya et al., 2013). The differentiation of pro-sensory cells into hair cells is triggered by their upregulation of the basic-helix-loop-helix (bHLH) transcription factor ATOH1, which is both necessary and sufficient for the production of hair cells (Bermingham et al., 1999; Zheng and Gao, 2000; Woods et al., 2004; Cai et al., 2013). SHH signaling opposes hair cell differentiation by maintaining the expression of bHLH transcriptional repressors and ATOH1 antagonists HEY1 and HEY2 within pro-sensory cells (Benito-Gonzalez and Doetzlhofer, 2014). Additional inhibitors of hair cell differentiation belong to the inhibitor of differentiation (ID) family (ID1-4). ID proteins are dominant negative regulators of bHLH transcription factors (Wang and Baker, 2015). Maintained by BMP signaling, ID1, ID2 and ID3 are thought to maintain pro-sensory cells in an undifferentiated state by interfering with ATOH1’s ability to activate its hair cell-specific target genes (Jones et al., 2006; Kamaid et al., 2010).

Much less is known about the signals and factors that promote ATOH1 expression/activity within pro-sensory cells and their role in auditory hair cell differentiation. Over-activation of Wnt/β-catenin signaling has been shown to increase *Atoh1* expression in differentiating cochlear explants and in the absence of Wnt/β-catenin signaling hair cells fail to form (Jacques et al., 2012; Shi et al., 2014; Munnamalai and Fekete, 2016). However, the pattern of Wnt–reporter activity, which at the onset of hair cell differentiation is high in the cochlear apex but low in the cochlear base, does not parallel the basal-to-apical wave of differentiation (Jacques et al., 2014). Interestingly, the *Inhba* gene, which encodes the Activin A subunit Inhibin βA (Barton et al., 1989), has been shown to be expressed in a basal-to-apical gradient within the differentiating auditory sensory epithelium (Son et al., 2015). Activins, which belong to the transforming growth factor (TGF)-β superfamily of cytokines, control a broad range of biological processes, including reproduction, embryonic axial specification, organogenesis and adult tissue homeostasis (reviewed in (Namwanje and Brown, 2016)). In the developing spinal cord, Activins and other TGF-β-related ligands are required in most dorsally located neuronal progenitors for *Atoh1* induction and their subsequent differentiation as D1A/B commissural neurons (Lee et al., 1998; Wine-Lee et al., 2004). The role of Activins in *Atoh1* regulation and hair cell differentiation are currently unknown.

Here, we identify Activin A signaling as a positive regulator of *Atoh1* gene expression and hair cell differentiation. We find that its graded pattern of activity within the auditory sensory epithelium times the basal-to-apical wave of hair cell differentiation. Furthermore, we provide evidence that Activin signaling informs a previously unidentified medial-to-lateral gradient of terminal mitosis that forces inner hair cell progenitors located at the medial edge of the sensory epithelium to withdraw from the cell cycle prior to outer hair cell progenitors.

## RESULTS

### The graded pattern of Activin A expression parallels auditory hair cell differentiation

The biological activity of Activin A and other Activins is limited by the secreted protein Follistatin (FST). Two FST molecules encircle the Inhibin β dimer, blocking both type I and type II receptor binding sites, thus preventing receptor binding and activation of its downstream signaling cascade (Thompson et al., 2005). Within the differentiating auditory sensory epithelium *Fst* and the Inhibin βA encoding gene *Inhba* are expressed in opposing gradients, with *Inhba* being highest expressed within the basal sensory epithelium and Fst being highest expressed apically (Son et al., 2015). To further characterize the pattern of *Inhba* and *Fst* expression and explore a potential correlation with hair cell differentiation, we carried out RNA in situ hybridization experiments using cochlear tissue stages E13.5-E15.5. *Sox2* expression was used to identify the pro-sensory/sensory domain, containing hair cell and supporting cell progenitors/precursors (Fig. 1J-L). Nascent hair cells were identified by their expression of *Atoh1* (Fig. 1 A-C). In mice, differentiation of basally located pro-sensory cells into hair cells starts at around embryonic stage E13.5-E14.0 (Fig.1 A). Paralleling *Atoh1* expression, *Inhba* expression was limited to the basal pro-sensory domain (Fig.1 D). In contrast, *Fst* was highly expressed within the lateral part of the pro-sensory domain throughout the cochlear apex and mid turn, but was only weakly expressed in the cochlear base (Fig. 1 G). At stages E14.5 and E15.5, as hair cell differentiation progressed towards the cochlear apex (Fig. 1 B, C), *Inhba* expression within the pro-sensory/sensory domain extended to the cochlear mid-turn (Fig.1 E, F). At the same time, *Fst* expression further regressed in the cochlear base and weakened in the cochlear mid-turn, but continued to be highly expressed in the undifferentiated cochlear apex (Fig. 1 H, I). As summarized in Fig.1 M, our analysis of *Inhba* and *Fst* expression indicates the existence of a basal-to-apical and medial-to-lateral gradient of Activin A signaling within the differentiating auditory sensory epithelium, which closely resembles the spatial and temporal pattern of hair cell differentiation, suggesting a link between Activin A signaling and hair cell differentiation.

**Figure 1:**
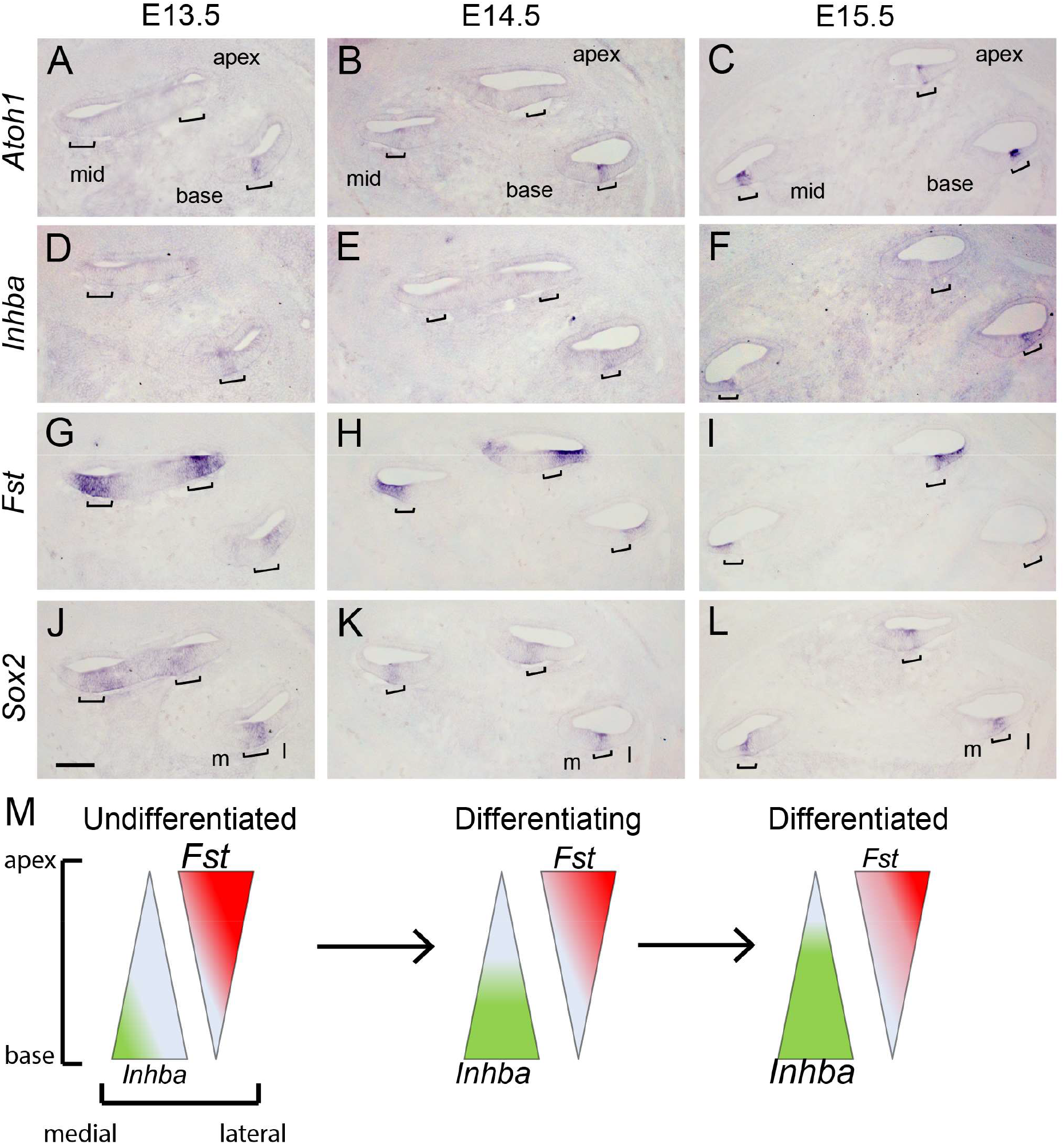
Activin A expression parallels auditory hair cell differentiation. A-L: In situ hybridization (ISH) was used to analyze the cochlear expression pattern of *Inhba* (D, E, F) and *Fst* (G, H, I) at the onset (E13.5) and during hair cell differentiation (E14.5 and E15.5). *Sox2* (J, K, L) expression was used to mark the pro-sensory/sensory domains, *Atoh1* (A, B, C) marks nascent hair cells. Brackets mark the pro-sensory/sensory domains with the cochlear duct. Abbreviations: m, medial; l, lateral. Scale bar, 100 μm. **M**: Schematics of longitudinal (apical–basal) and radial (medial-lateral) expression gradients of *Inhba* and *Fst* within the auditory sensory epithelium.

### Activin signaling promotes auditory hair cell differentiation

Activins play a central role in coordinating growth and differentiation in sensory and neuronal tissues, including the spinal cord, retina and olfactory epithelium (Liem et al., 1997; Davis et al., 2000; Gokoffski et al., 2011). To determine the role of Activin A in cochlear hair cell differentiation, we exposed undifferentiated cochlear tissue (stage E13.5) to recombinant Activin A (500 ng/ml) for 24 hours (Fig. 2 A). To monitor the dynamics of hair cell differentiation, we made use of transgenic mice that carried the enhanced green fluorescent protein (EGFP) transgene under the control of the *Atoh1* enhancer (Lumpkin et al., 2003). Hair cell differentiation follows a steep basal-to-apical gradient in which hair cells located within the cochlear mid-basal segment differentiate prior to more apical located hair cells. In addition, a shallower, medial-to-lateral gradient exists, with inner hair cells differentiating prior to their neighboring more laterally located outer hair cells (Chen et al., 2002). Whereas control cultures showed no or only weak Atoh1-reporter activity (EGFP) within the developing cochlear duct after 12 hours of culture (Fig. 2 B), Activin A-treated cochlear cultures already contained a narrow strip of EGFP positive inner hair cells (Fig. 2 C). 6 hours later, a broad strip of EGFP positive cells representing inner and outer hair cells was evident in Activin A-treated cochlear explants (Fig. 2 E), whereas control cochlear explants contained only few scattered EGFP positive inner hair cells (Fig. 2 D). Furthermore, at all stages examined, the band of EGFP positive cells extended further apically in Activin A treated cochlear explants compared to control (Fig. 2 B-G, H). Taken together our findings indicate that Activin A acts a differentiation signaling for auditory hair cells.

**Figure 2:**
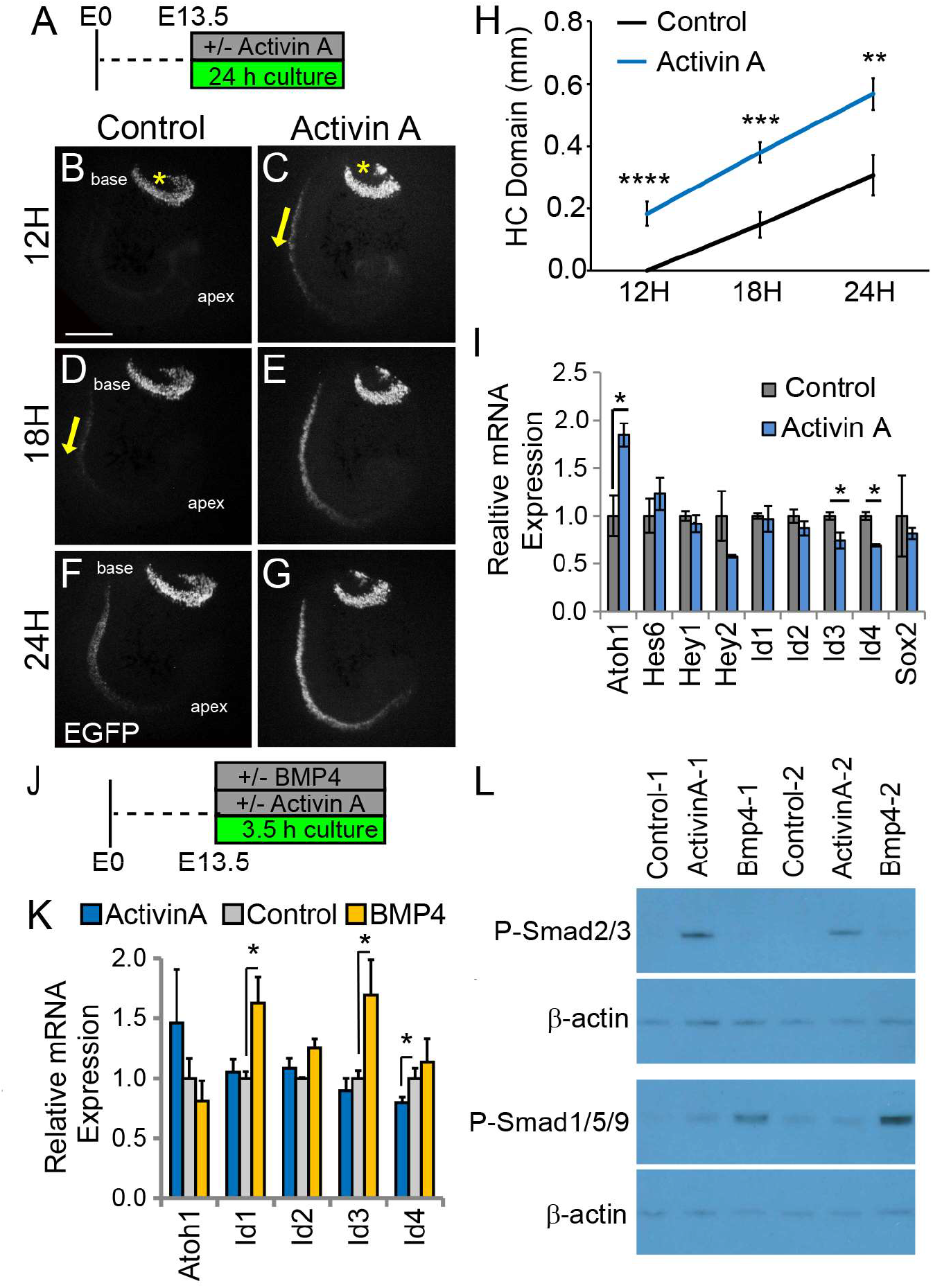
Activin A promotes auditory hair cell differentiation. **A**: Experimental design for B-I. Stage E13.5 wild type cochlear explants were cultured with or without Activin A (final conc. 500 ng/ml) for 24 hours. **B-G** Atoh1/nEGFP reporter expression (EGFP, gray) was used to monitor and analyze hair cell differentiation in Activin treated (C, E, G) and control (B, D, F) cochlear explants. Asterisks marks vestibular sacculus that contains Atoh1-positive hair cells. Yellow arrows mark the onset of hair cell differentiation within the cochlea. Scale bar, 100 μm. **H**: Quantification of extent of hair cell differentiation in control versus Activin treated cochlear cultures (see B-G). Data expressed as mean ± SEM (n = 5-8 cochlear explants per group, **p < 0.01, ***p < 0.001, ****p < 0.0001). **I**: Transcript levels of pro-sensory genes (*Id1-4, Hey1, Hey2* and Sox2) and hair cell-specific genes (*Atoh1, Hes6*) were analyzed in enzymatically purified cochlear epithelia. 3-4 cochlear epithelia were pooled per sample. Data are mean ± SEM (n=3 independent experiments, *p < 0.05). J: Experimental design for K, L. Stage E13.5 wild type cochlear epithelia were cultured with or without Activin A (final conc. 200 ng/ml) or BMP4 (final conc. 100ng/ml) for 3.5 hours. **K**: RT-qPCR analysis reveals differential response to Activin and BMP treatment. Individual cochlear epithelia were analyzed. Data are mean ± SEM, n=4 biological replicates, *p < 0.05. **L**: Activin A induces Smad2/3 phosphorylation in cochlear epithelial cells. Western blot analysis was used to establish P-Smad2/3 and P-Smad1/5/9 protein levels in individual cochlear epithelial extracts after 3.5 hour exposure to Activin A or BMP4. Beta-actin was used as loading control.

To identify target genes of Activin signaling in the developing cochlea, we cultured E13.5 cochlear explants with or without Activin A for 24 hours, after which we enzymatically purified the corresponding cochlear epithelia and performed RT-qPCR. Our analysis focused on key regulators of hair cell differentiation, which included the pro-hair cell factor ATOH1, the sensory specification factor *SOX2* (Neves et al., 2012; Kempfle et al., 2016), the hair cell fate repressors HEY1, HEY2 (Benito-Gonzalez and Doetzlhofer, 2014), as well as members of the inhibitor of differentiation (ID) family (ID1-4). We found that Activin A treatment did not significantly alter mRNA abundance of *Hey1, Hey2, *Sox2*, Id1* and *Id2* expression. However, we found that in response to Activin treatment *Atoh1* transcript was significantly increased by more than 2-fold and that *Id3* and *Id4* mRNA abundance was significantly reduced (Fig. 2 I).

To untangle direct and indirect effects of Activin signaling on *Id3, Id4* and *Atoh1* gene expression we shortened the culture period to 3.5 hours and eliminated non-epithelial tissue from our cultures. In addition, we included a brief BMP4 treatment as positive control, allowing us to compare the effects of Activin A and BMP4 on pro-sensory/sensory gene expression.

BMP signaling is known to positively regulate *Id1-3* expression in inner ear pro-sensory cells (Kamaid et al., 2010; Ohyama et al., 2010). Consistent with these previous reports, 3.5 hour BMP4 treatment of E13.5 cochlear epithelia led to an increase in *Id1-3* expression, which was significant for *Id1* and *Id3*, but showed no effect on *Id4* expression compared to control (Fig. 2 K). In contrast, 3.5 hour Activin A treatment of E13.5 cochlear epithelia had no effect on *Id1-3* expression but resulted in a modest, but significant decrease in *Id4* expression compared to control (Fig. 2 K). The divergent transcriptional responses to Activin A and BMP4 are likely the consequence of differences in type I receptor and R-SMAD utilization. Activins commonly utilize the type I receptor ALK4 (ACVR1B), which signals through SMAD2 and SMAD3, whereas BMPs activate the type I receptors ALK3 (BMPR1A) and ALK6 (BMPR1B), which signal through SMAD1, SMAD5 and SMAD9 activation (reviewed in (Miyazawa et al., 2002)). Indeed, we found that a 3.5-hour exposure of E13.5 cochlear epithelia to Activin A selectively stimulated the phosphorylation of SMAD2/3, whereas a 3.5-hour treatment with BMP4 resulted in the phosphorylation of SMAD1/5/9 (Fig. 2 J, K). In contrast to the 24-hour Activin A treatment, the 3.5-hour Activin A treatment failed to significantly increase *Atoh1* expression, suggesting that Activin A promotes the expression of *Atoh1* through indirect mechanisms.

### FST inhibits auditory hair cell differentiation

To characterize the function of FST in cochlear development we made use of a recently developed doxycycline (dox) inducible transgenic mouse line, in which a cassette encoding the human FST-288 isoform is under the control of a tetracycline-responsive promoter element (tetO) (Lee and McPherron, 2001; Roby et al., 2012) (Fig. 3 A). In the presence of a ubiquitously expressed reverse tetracycline-controlled trans-activator (rtTA), dox administration allows for robust induction of the human (h) FST transgene (Fig. 3 A, B, G). To assess the role of FST in cochlear hair cell differentiation, timed pregnant females received dox beginning at E11.5 and double transgenic FST over-expressing embryos (R26-FST) and single transgenic littermates lacking the R26-M2rtTA transgene (control), were harvested three days later (E14.5). Inclusion of the *Atoh1*-reporter transgene (*Atoh1/nEGFP*) allowed for the ready analysis of hair cell differentiation at the time of isolation. While Atoh1-reporter positive inner hair cells were detectable in the cochlear base of stage E14.5 control cochlear tissue (Fig. 3 C, E), stage E14.5 FST overexpressing cochlear tissue lacked cochlear hair cells (Fig. 3 D, F), indicating an inhibitory function for FST in auditory hair cell differentiation.

**Figure 3:**
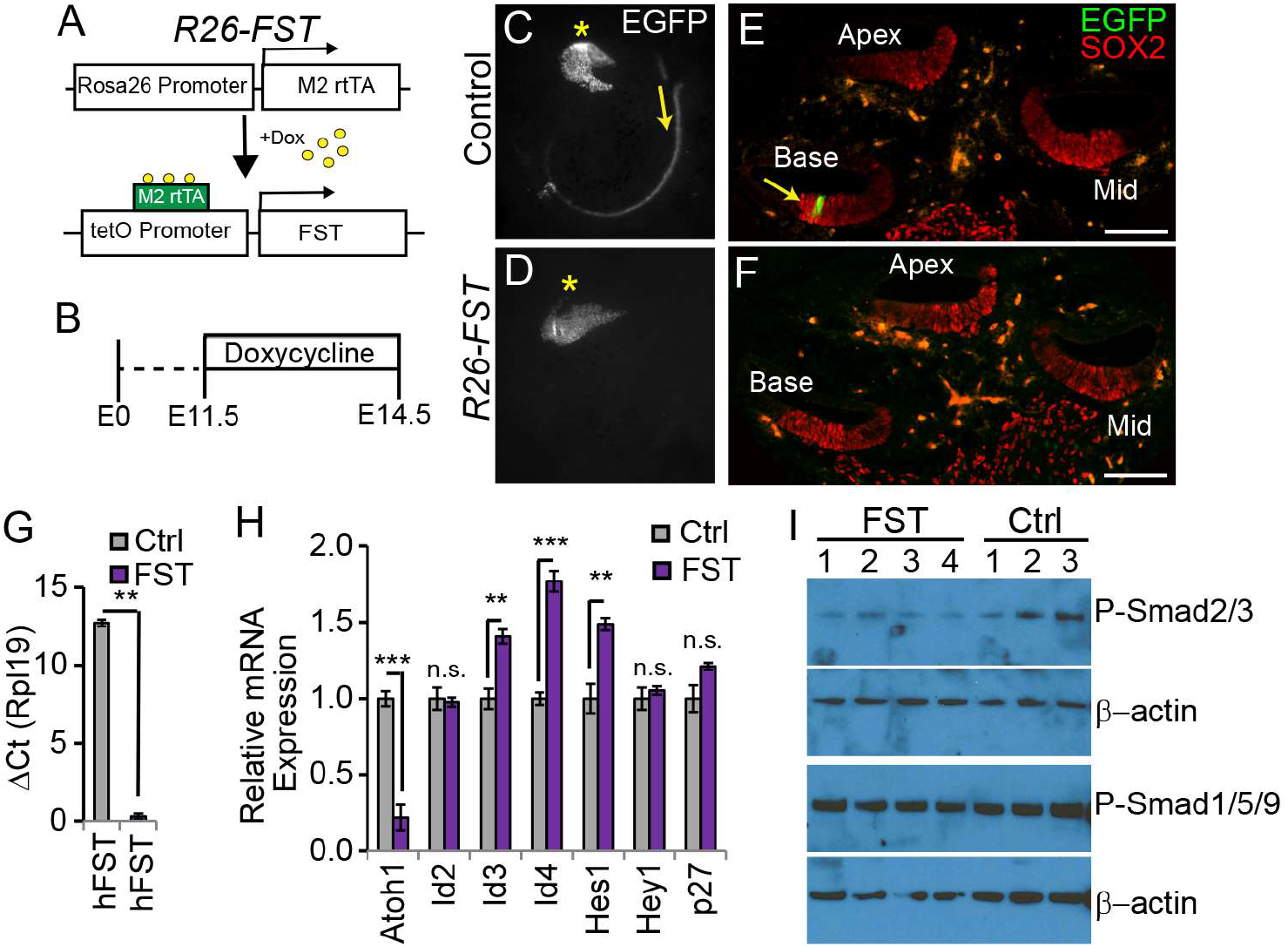
FST inhibits auditory hair cell differentiation. **A**: Inducible FST transgenic mouse model. In the presence of doxycycline (dox) double transgenic animals (R26-M2rtTA and tetO-hFST) express human FST under the control of the R26 promoter (R26-FST). Non-transgenic littermates and littermates that carry only one of the transgenes were used as experimental controls (Ctrl). **B**: Experimental strategy for C-H. Timed mated pregnant dams received dox at stage E11.5 and FST transgenic (R26-FST) animals and control littermates (Ctrl) were harvested at E14.5. **C, D**: Low power fluorescent images of native Atoh1/nEGFP reporter expression (EGFP, grey) of FST overexpressing (D) and control cochlear epithelia. Scale bar 100 μm. **E, F**: Confocal images FST overexpressing (F) and control cochlear cross sections (E). Native Atoh1/nEGFP (EGFP, green) marks hair cells (yellow arrow); SOX2 staining (red) marks the sensory domain. Scale bar 100 μm. **G**: Human (h) *FST* transgene expression in control (Ctrl) and *R26-FST* transgenic (FST) cochlear epithelia. Plotted is their relative quantity (ΔCT) compared to the reference gene Rpl19. Data expressed as mean ± SEM (n = 4-5 animals per group, **p<0.01). **H**: RT-qPCR-based analysis of gene expression in FST overexpressing (FST) and control cochlear epithelia (Ctrl). Data are mean ± SEM (n = 4-5 animals per group,* p< 0.05, **p < 0.01, ***p< 0.001). **I**: Western-blot based analysis of P-SMAD2/3 and P-SMAD1/5/9 protein levels in *FST* overexpressing (R26-FST: 1-4) and control (control: 1-3) cochlear epithelia. Timed mated pregnant dams received dox at starting at E12.5 and cochlear epithelia were isolated two days later from stage E14.5 FST transgenic (R26-FST) animals and control littermates (Ctrl). Beta-actin was used as loading control.

To determine how FST overexpression interferes with hair cell differentiation we again administered dox at E11.5, isolated FST transgenic and cochlear epithelia from stage E14.5 FST overexpressing and control embryos and analyze their pro-sensory/ sensory gene expression using RT-qPCR. We found that FST overexpression significantly reduced *Atoh1* expression and significantly increased *Hes1, Id3* and *Id4* expression, but did not significantly alter *p27/Kip1, Id2* or *Hey1* expression (Fig. 3 H). Activins as well as the Activin–type ligands Gdf11 and myostatin are high affinity binding partners for FST (Harrington et al., 2006). In addition, FST has been shown to bind at low affinity to BMP4 and BMP7 and modulate their activity (Iemura et al., 1998; Amthor et al., 2002). These two related BMP ligands are abundantly expressed in the developing cochlea and are thought to function in the specification and patterning of the pro-sensory domain (Ohyama et al., 2010). To determine whether FST overexpression inhibits BMP signaling in the developing cochlea, we administered dox at E12.5, and two days analyzed P-SMAD2/3 and P-SMAD1/5/9 proteins in acutely isolated control and FST overexpressing cochlea epithelia. Our western blot analysis revealed that P-SMAD2/3 protein levels were reduced in FST overexpressing cochlear epithelia compared to control cochlear epithelia. However, FST overexpression had little to no effect on P-SMAD1/5/9 protein levels, indicating that FST overexpression in the developing cochlea selectively disrupts Activin-type signaling (Fig. 3 E).

To determine whether exogenous Activin A can rescue the FST mediated delay in hair cell differentiation, we monitored *Atoh1-reporter* expression in cultures of E13.5 FST overexpressing (R26-FST) and single transgenic wild type cochlear tissue treated with or without Activin A (Fig. 4 A). Consistent with our earlier findings, we observed that hair cell differentiation occurred significantly earlier in Activin A-treated wild type cochlear explants compared to untreated wild type cochlear explants (Fig. 4 C, B, N). Conversely, the onset of hair cell differentiation was significantly delayed in untreated FST overexpressing cochlear explants compared to untreated wild type cochlear explants (Fig. 4 F, H, N). However, these defects were almost completely abolished when Activin A treatment and FST overexpression were combined and no significant differences in the onset (Fig. 4 F, I, N) or in the progression of hair cell differentiation was observed between FST overexpressing cochlear explants treated with Activin A and wild type untreated cochlear explants (Fig. 4 J, M, N). Taken together, our data suggests that FST antagonizes hair cell differentiation in an Activin A-dependent manner.

**Figure 4:**
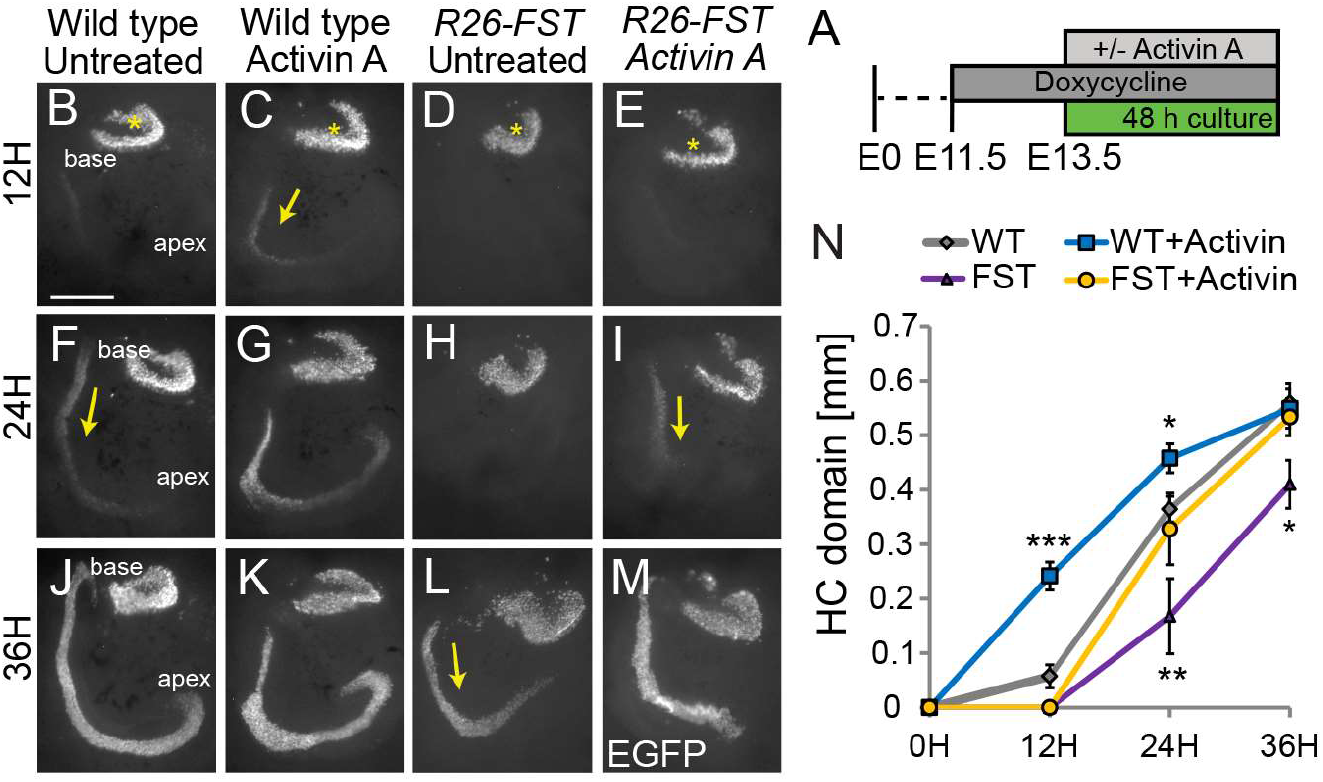
Exogenous Activin A rescues the FST induced delay in auditory hair cell differentiation. **A**: Experimental design for B-N. Dox was administered to timed pregnant dams starting at E11.5. At E13.5, cochlear tissue from FST overexpressing embryos and control littermates were cultured for 48 hours with or without Activin A (500 ng/ml). **B-M**: Atoh1/nEGFP reporter expression (EGFP, grey) marks nascent hair cells. Asterisks indicate hair cells within vestibular sacculus. Yellow arrows mark nascent cochlear hair cells. Scale bar, 100μm. **N**: Length of Atoh1/nEGFP positive sensory epithelium was used to quantify the extent of hair cell differentiation in control and FST overexpressing cochlear explants cultured with and without Activin A. Data expressed as mean ± SEM (n = 5-18 cochlear explants per group, p ≤ 0.05, **p < 0.01, ***p < 0.001). Two independent experiments were conducted.

### FST promotes pro-sensory cell proliferation

In the mammalian cochlea, cell cycle exit of pro-sensory cells occurs in a highly synchronized manner. Pro-sensory cells located in the cochlear apex withdraw from the cell cycle first, followed by more basally located pro-sensory cells (Ruben, 1967; Lee et al., 2006). To determine whether FST antagonizes pro-sensory cell cycle withdrawal, we injected timed pregnant dams with the thymidine analog EdU at stage E13.5, which in mice corresponds to the peak of pro-sensory cell cycle withdrawal, and analyzed hair cell and supporting cell-specific EdU incorporation in FST overexpressing embryos and control littermates five days later (stage E18.5). Hair cells were identified by their native EGFP (Atoh1/nEGFP) expression as well as by immuno-staining for the hair cell-specific protein myosinVIIa (Myo7a); supporting cells were identified by SOX2 immuno-staining. Classification of cell subtypes was based on their location within the sensory epithelium. As expected we found that basally located hair cells and their surrounding supporting cells incorporated EdU at high frequency in both control (Fig. 5 A) and FST overexpressing cochlear tissue (Fig. 5 D). However, whereas in control cochlear tissue supporting cells and hair cells located further apically (mid and apex) showed little to no EdU incorporation (Fig. 5 B, C), their counterparts in FST overexpressing cochleae exhibited robust EdU incorporation (Fig. 5 E, F), indicating that FST overexpression in the developing cochlea delays pro-sensory cell cycle withdrawal. The number and percentage of EdU labeled inner and outer hair cells was significantly lower in the apex than in the mid-turn of FST overexpressing cochleae, indicating that FST overexpression prolongs pro-sensory cell proliferation without disrupting the apical-to-basal gradient of pro-sensory cell cycle withdrawal (Fig. 5 G, H). Our analysis did however find that FST overexpression alters a previously unrecognized medial-to-lateral gradient of terminal mitosis within the auditory sensory epithelium. In the basal segment of the control and FST overexpressing auditory sensory epithelia, in which, at the time of EdU administration pro-sensory cells had been actively dividing, the percentage of EdU positive hair cells that were inner hair cells correlated with their relative abundance (Fig. 5 I, base). However, in the mid-turn of control cochleae, where pro-sensory cells had been in the process of withdrawing from the cell cycle, no EdU positive inner hair cells were detected and 100% of the EdU labeled hair cells were outer hair cells, suggesting that inner hair cell progenitors had exited the cell cycle prior to the more laterally located outer hair cell progenitors. However, in FST overexpressing cochleae this radial gradient of terminal mitosis appeared to be lost (Fig. 5 I, mid).

**Figure 5:**
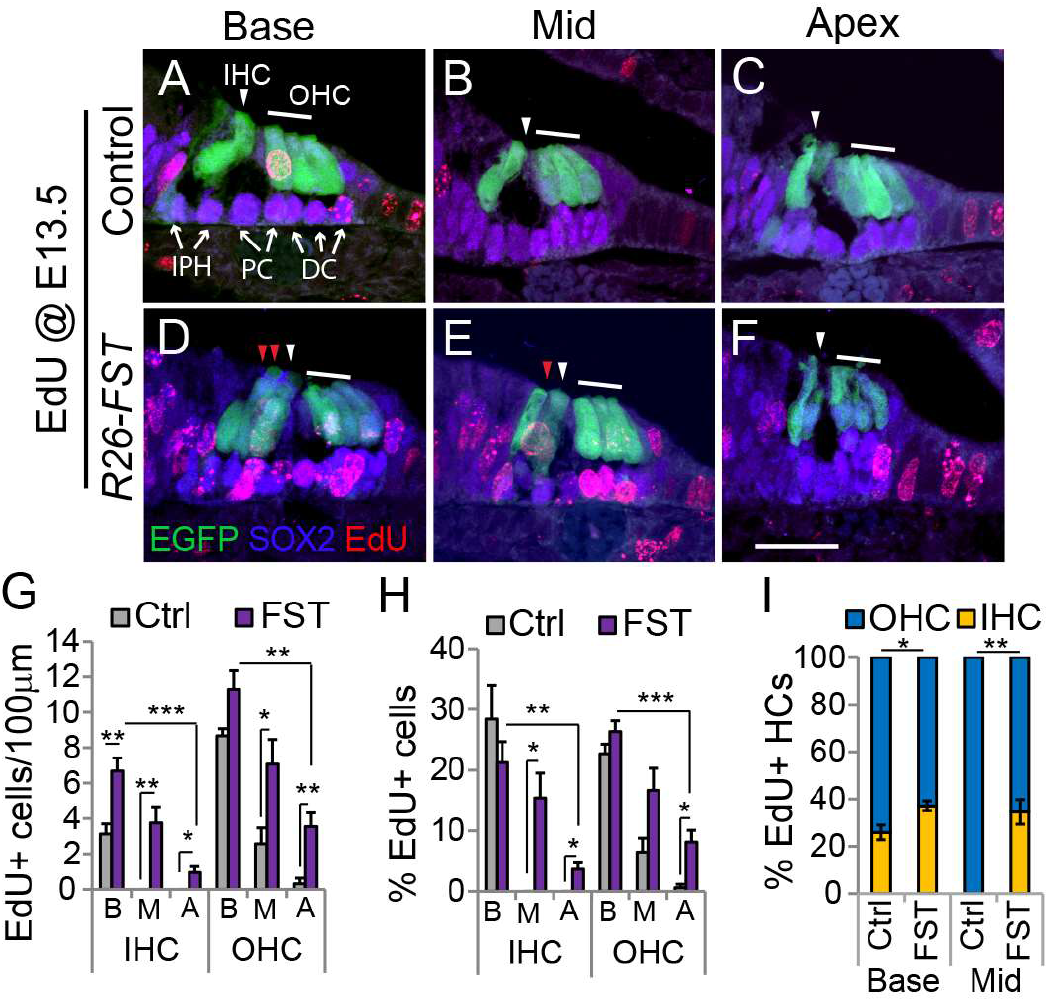
FST overexpression delays pro-sensory cell cycle exit. To induce *FST* transgene expression, timed mated pregnant dames received dox beginning at E11.5. A single EdU pulse was administered at E13.5 and EdU incorporation (red) in hair cells and supporting cells was analyzed at stage E18.5. **A-F**: Confocal images of cross-sections through the base, mid and apical turn of control (A-C) and FST overexpressing (D-F) cochlear tissue. Atoh1/nEGFP transgene expression (green) marks inner hair cells (IHC, white arrowhead) and outer hair cells (OHC, white bar). SOX2 immunostaining (magenta) marks supporting cells including inner phalangeal cells (IPH), pillar cells (PC) and Deiters’ cells (DC) marked by white arrows. Ectopic inner hair cells are marked by red arrowheads. Scale bar, 50 μm. **G-I**: Quantification of EdU positive hair cells within the base, mid and apex of control (Ctrl, grey bars) and FST overexpressing (FST, purple bars) cochlear whole mounts. Graphed are the number of EdU positive inner and outer hair cells per 100 μm (G), the percentage of inner and outer hair cells that are EdU positive (H) and the percentage of EdU positive hair cells that are inner versus outer hair cells (I). Abbreviations: IHC, inner hair cells; OHC, outer hair cells; B, base; M, mid; A, apex. Data expressed as mean ± SEM (n =4 animals per group, *p ≤ 0.05, **p < 0.01, ***p < 0.001).

To further characterize the effects of FST on inner hair cell progenitor proliferation, we administered EdU from E14.5 to E17.5 and analyzed EdU incorporation in hair cells and supporting cells at E18.5. In control cochlear tissue, only few, most basally located hair cells and supporting cells were labeled with EdU (Fig. 6 A-C, G, H), indicating that the majority of pro-sensory cells had withdrawn from the cell cycle prior to stage E14.5. In contrast, robust EdU labeling of supporting cells and hair cells was observed throughout the base and mid-turn of FST overexpressing cochleae (Fig. 6 D-F, G, H). Again, in control tissue inner hair cells that were inner hair cells was increased in FST overexpressing cochlear tissue compared to control Consistent with the existence of a medial to lateral gradient of cell cycle withdrawal, in the base of control cochlear tissue, 94 % of all EdU labeled hair cells were outer hair cells and only 6% of all EdU labeled hair cells were inner hair cells. In contrast, in the base of FST overexpressing cochlear tissue 51% of all EdU labeled hair cells were inner hair cells (Fig. 6 I). In the apex of FST overexpressing cochleae, inner hair cells and inner phalangeal cells were the only cell types that were labeled with EdU (Fig. 6 F, G, H). In summary, our findings indicate that FST acts as a mitogen for pro-sensory cells, in particular inner hair cell progenitors and identify FST as a key regulator of a newly uncovered radial gradient of pro-sensory cell cycle withdrawal.

**Figure 6:**
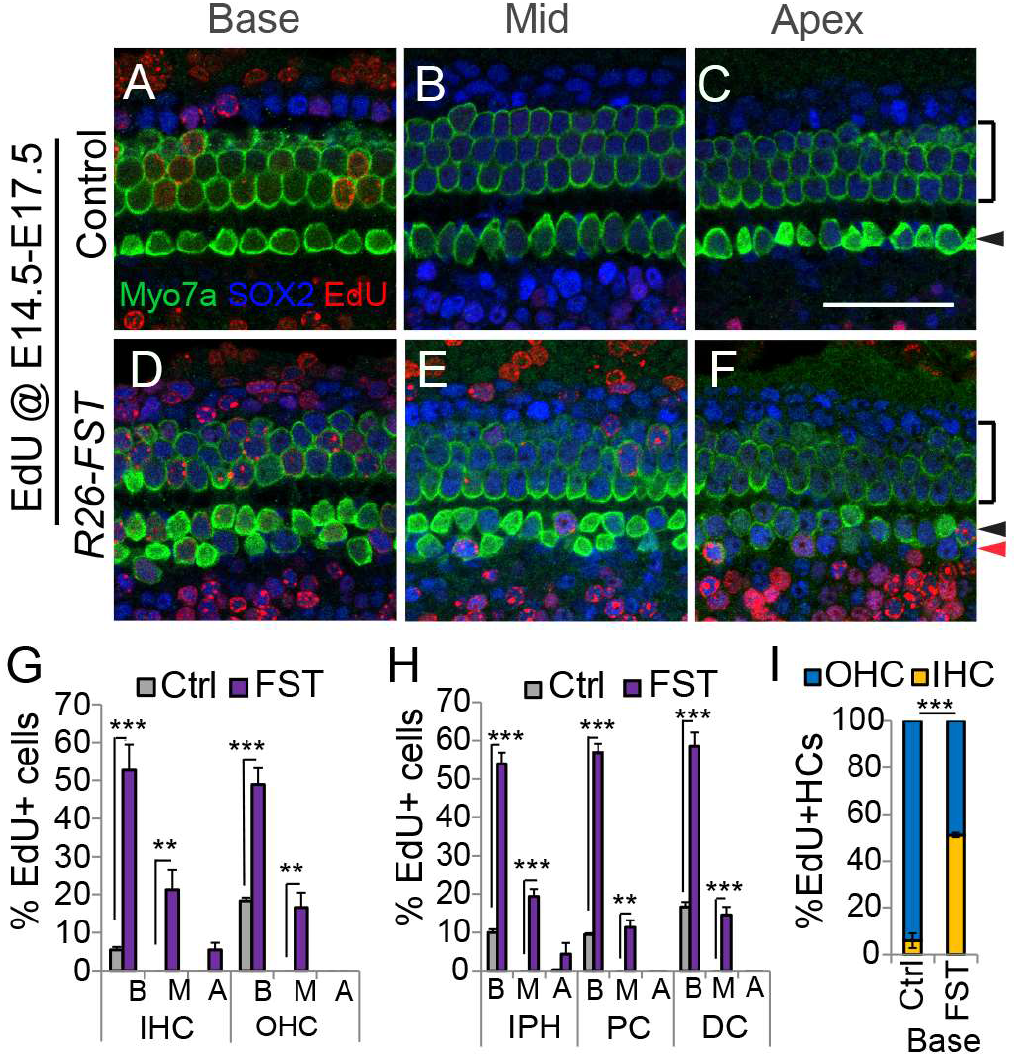
FST overexpression increases inner hair cell progenitor proliferation. To induce *FST* transgene expression, timed mated pregnant dames received dox beginning at E11.5. Timed mated pregnant dams received two pulses of EdU daily starting from E14.5 to E17.5 and EdU incorporation (red) in hair cells and supporting cells was analyzed at stage E18.5. **A-F**: Confocal images of representative fields of basal, mid and apical segments of control (A-C) and FST overexpressing (D-F) auditory sensory epithelia. Myo7a immuno-staining (green) marks inner (black arrowhead), outer (black bar) and ectopic inner hair cells (red arrow head). Nuclear SOX2 immuno-staining (blue) marks surrounding supporting cells and less mature hair cells. Scale bar, 50 μm. **G-I**: Quantification of EdU positive hair cells and supporting cells within the base, mid and apex of control (Ctrl, grey bars) and FST overexpressing (FST, purple bars) cochlear whole mounts. Graphed are the percentage of hair cell (G) and supporting cell sub-types (H) that are EdU positive and the percentage of EdU positive hair cells that are inner versus outer hair cells (I). Abbreviations: IHC, inner hair cells; OHC, outer hair cells; IPH, inner phalangeal cells; PC, pillar cell; DC, Deiters’ cell; B, base; M, mid; A, apex. Data expressed as mean ± SEM (n = 3 animals per group *p ≤ 0.05, **p < 0.01, ***p < 0.001).

### Activin signaling controls the differentiation and cellular patterning of hair cells

As typical for wild type cochlear tissue, cochlear tissue from stage E18.5 control embryos contained a single row of inner hair cells and 3 rows of outer hair cells (Fig. 7 A, C, E). However, likely a consequence of continued proliferation of inner hair cell progenitors, cochlear tissue from FST overexpressing embryos contained 2-3 rows of inner hair cells. The ectopic inner hair cell phenotype was most severe in the base of the cochlea; inner hair cell density was increased nearly 2.5-fold compared to controls and was accompanied by an increase in surrounding inner phalangeal cells (Fig. 7 B, D, G, H). The density and cellular patterning of outer hair cells was largely unchanged in FST overexpressing cochleae except for in the cochlear base. The base of FST overexpressing cochleae contained stretches of sensory epithelium in which the 3^rd^ row of outer hair cells and the 3^rd^ row of accompanying Deiters’ cells were missing, resulting in reduced outer hair cell and Deiters’ cell densities (Fig. 7 D, G, H). Furthermore, differentiation/maturation of the auditory sensory epithelium was severely delayed in response to FST overexpression. At stage E18.5, the basal-to-apical wave of differentiation has reached the cochlear apex. The initially multilayered auditory sensory epithelium is largely thinned to a two-layered sensory epithelium and is near its final length. Moreover, hair cells, including hair cells located in the cochlear apex, have formed actin-rich apical protrusions, so called stereocilia (see Fig. 7 C, E). Consistent with a severe delay in pro-sensory cell differentiation, hair cells in FST overexpressing cochlear tissue had less developed stereocilia than their wild type counterparts, and apical hair cells lacked stereocilia completely (Fig. 7 C-F). Moreover, our analysis revealed that the auditory sensory epithelia of FST overexpressing cochleae consisted of more than two epithelial layers (see Fig. 5 A-F) and were 30% shorter compared to control sensory epithelia obtained from littermates (dox E11.5, harvest E18.5: control = 5.14± 0.04 mm, R26-FST =3.65 ± 0.13 mm, n=5, p=0.0002).

**Figure 7:**
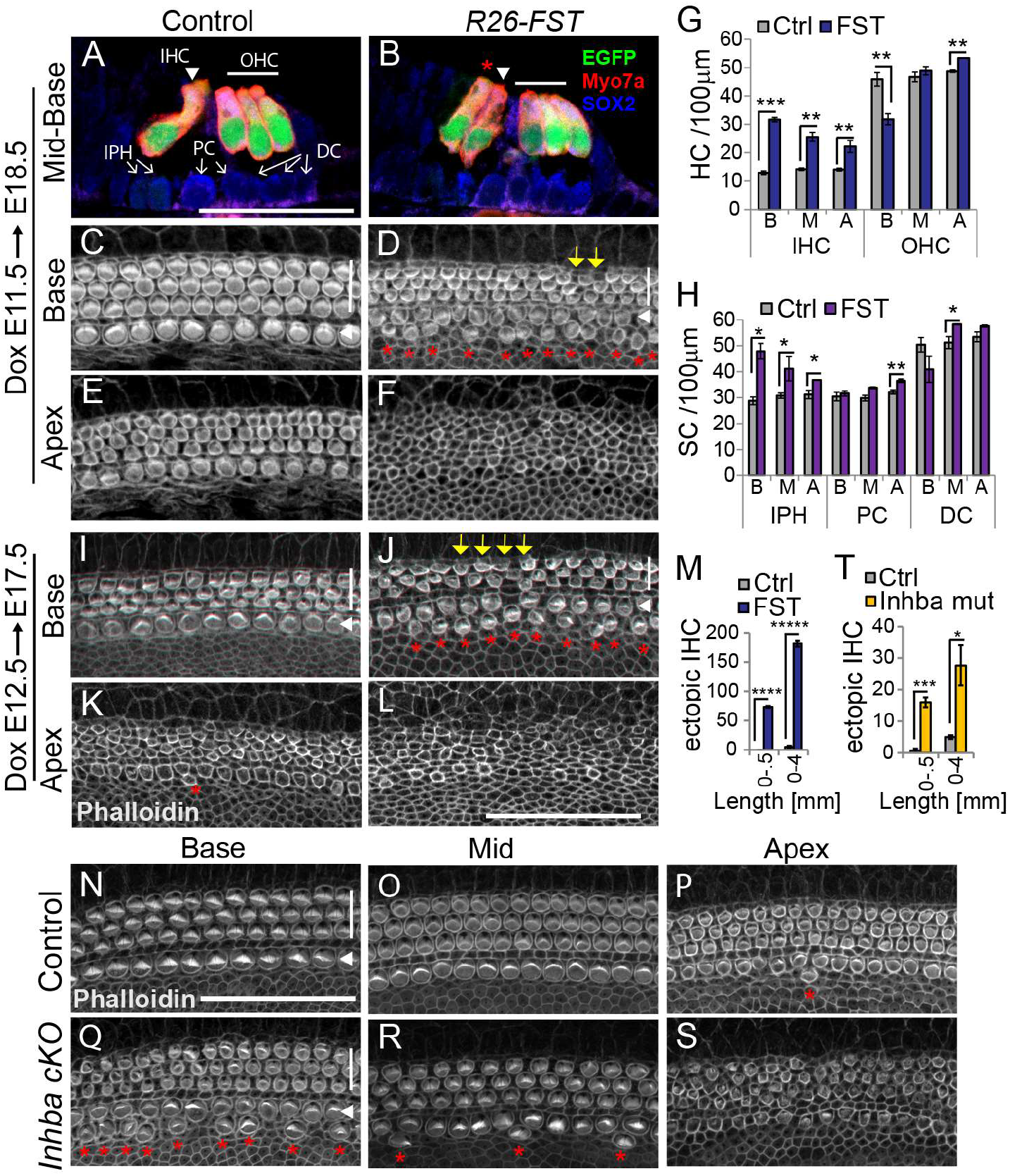
Disruption of Activin signaling increases inner hair cell formation and delays hair cell differentiation. **A-T**: FST transgenic (R26-FST) embryos and their control (wild type or single transgenic) littermates were exposed to dox starting at E11.5 until tissue harvest at E18.5 (A-H) or starting at E11.5 until tissue harvest at E17.5 (I-T). **A-B: FST overexpression results in ectopic inner hair cell formation**. Confocal images of cross-sections through the midbase of control (A) and *FST* overexpressing (B) cochleae. Atoh1/nEGFP (green) and Myo7a (red) labels inner (white arrowhead), outer (white bar) and ectopic inner hair cells (red asterisk). SOX2 (blue) labels supporting cells including inner phalangeal cells (IPH), pillar cells (PC) and Deiters’ cells (DC) indicated by white arrows. Scale bar 50μm. **C-L: FST overexpression delays hair cell differentiation/maturation**. Confocal images of the apical surface of hair cells located at the base (C, D, I, J) and apex (E, F, K, L) of *FST* over-expressing (D, F, J, L) and control (C, E, I, K) cochlear whole mount preparations. Phalloidin labels actin-rich stereocilia of inner (white arrowhead) and outer hair cells (white bar). Red asterisks mark ectopic inner hair cells. Yellow arrows mark location of missing outer hair cells. Scale bar 50μm **G-H**: Quantification of hair cells (G) and supporting cell (H) density in the base, mid and apex of control (Ctrl, grey bars) and FST overexpressing (FST, purple bars) cochleae. Abbreviations: IHC, inner hair cells; OHC, outer hair cells; IPH, inner phalangeal cells; PC, pillar cell; DC; Deiters’ cell, B, base, M, mid, A, apex. Data expressed as mean ± SEM (n = 3 animals per group, *p ≤ 0.05, **p<0.01, ***p<0.001). **M**: Graphed are the number of ectopic inner hair cells (IHC) within the most basal segment (0-.5 mm) and within the entire length (0 – 4mm) of control (Ctrl, grey) and FST overexpressing (FST, purple) cochleae. Data expressed as mean ± SEM (n = 3 animals per group, ****p < 0.0001, ***** p < 0.00001). **N-T: Loss of Inhba delays hair cell maturation and causes a mild overproduction of inner hair cells**. Shown are confocal images of the apical surface of hair cells located in the base (N, Q), mid (O, R) and apex (P, S) of stage P0 *Inhba* mutant (*Inhba* cKO) (Q-S) and control (Ctrl) cochlear whole mount preparations (N-P). Phalloidin labels actin-rich stereocilia of inner (white arrowhead) and outer hair cells (white bar). Red asterisks mark ectopic inner hair cells. Scale bar 50 μm. T: Graphed are the number of ectopic inner hair cells (IHC) within the most basal segment (0–.5 mm) and within the entire length (0 – 4mm) of control (Ctrl, grey) and *Inhba* mutant (*Inhba* mut, orange) cochleae. Data expressed as mean ± SEM (n = 3 animals per group, *p ≤ 0.05, ***p < 0.001).

FST induction one day later at~ E13.0 (dox E12.5), had no effect on the length of the auditory sensory epithelium (dox E12.5, harvest E17.5: control = 4.13± 0.17 mm, R36-FST =3.99 ± 0.12 mm, n=3, p=0.5425), but we continued to observe ectopic inner hair cells in FST overexpressing cochleae, with the highest number of ectopic hair cells found in the basal portion of the auditory sensory epithelium, as well as short stretches of sensory epithelium with only 2 rows of outer hair cells (Fig. 7 J, M). Furthermore, we found that hair cell stereocilia in FST overexpressing cochlea had a less mature phenotype compared to their counterparts in control cochlear tissue, indicating that FST induction as late as ~E13.0 causes a delay in hair cells differentiation/maturation (Fig. 7 I-L).

The formation of inner hair cells is regulated by a complex interplay of various signaling pathways including Notch, Wnt and BMP signaling (reviewed in (Groves and Fekete, 2012)). In particular, a recent study revealed that inhibition of BMP signaling using dorsomorphin results in the overproduction of inner hair cells in cochlear explants (Munnamalai and Fekete, 2016).

To determine whether FST influences inner hair cell formation in an Activin A-dependent manner, we analyzed hair cell patterning in Activin A (Inhba) mutant mice. To selectively ablate *Inhba* gene function in the developing inner ear, we intercrossed *Inhba* floxed mice (*Inhba* fl/fl), in which exon 2 of the *Inhba* gene is flanked by LoxP sites (Pangas et al., 2007), with inner ear-specific *Pax2-Cre* transgenic mice (Ohyama and Groves, 2004). Conditional knockout (cKO) of Inhba gene function did not alter overall cochlear morphology, nor did it alter the length of the sensory epithelium compared to control (*Inhba* fl/fl) (P0: control = 4.99 ± 0.12 mm; *Inhba* cKO = 4. 82 ± 0.01 mm; n=2, p-value=0.4043). However, our analysis revealed defects in inner hair cell patterning in the absence of *Inhba* that were qualitatively similar to inner hair cell patterning defects observed in response to FST overexpression (dox E12.5). Cochlear tissue from control littermates (*Inhba* fl/fl) contained the normal compliment of one row of inner hair cells and three rows of outer hair cells (Fig. 7 N-P). In contrast, *Inhba* cKO (*Pax2-Cre; Inhba* fl/fl) cochlear tissue contained ectopic inner hair cells, with much of the ectopic inner hair cells residing within the most basal segment (Fig. 7 Q-S, T). Furthermore, consistent with a delayed onset of hair cell differentiation, hair cell stereocilia in the base, mid and apex of *Inhba* mutant cochlear tissue had a less mature phenotype compared to control cochlear tissue (Fig. 5 N-S). In summary, these findings indicate that Activin A-mediated signaling is critical for proper inner hair cell formation as well as hair cell differentiation/maturation *in vivo*.

## DISCUSSION

Signaling gradients play a fundamental role in controlling growth and differentiation during embryonic development. Here, we show that Activin A acts as differentiation signal for auditory hair cells and provide evidence that its graded activity times the longitudinal gradient of differentiation within the auditory sensory epithelium. Furthermore, we identify FST as an antagonist of Activin signaling and show that FST overexpression delays pro-sensory cell cycle exit and differentiation. Finally, we provide evidence that Activin signaling regulates a previously unrecognized radial gradient of pro-sensory cell cycle withdrawal, limiting the number of inner hair cells being produced.

### Activin A functions as a pro-differentiation signal in the mammalian cochlea

How does Activin signaling promote auditory hair cell differentiation? Our findings suggest that Activin A facilitates hair cell differentiation through boosting the expression of the pro-hair cell transcription factor ATOH1 within pro-sensory cells. The time required to observe robust increase in *Atoh1* expression in response to exogenous Activin A suggests that Activin A promotes *Atoh1* expression through indirect mechanisms. In particular, we found that Activin signaling reduced the expression of bHLH antagonist ID3 and ID4 within pro-sensory cells. ID proteins function as dominant-negative regulators of bHLH transcription factors by binding to bHLH E-proteins, which are required for bHLH transcription factors such as ATOH1 to form functional hetero-dimers capable of binding DNA (Benezra et al., 1990; Wang and Baker, 2015). Consistent with acting as an ATOH1 antagonist, ID3 overexpression in the developing inner ear inhibits hair cell formation in both auditory and vestibular sensory organs (Jones et al., 2006; Kamaid et al., 2010). The role of ID4 in inner ear development has yet to be established.

It should be noted that *Atoh1* expression and subsequent hair cell differentiation is not disrupted, but rather delayed in response to FST overexpression or *Inhba* ablation, suggesting the existence of Activin-independent mechanisms capable of activating *Atoh1* expression in pro-sensory cells. A likely candidate is the Wnt/β-catenin signaling pathway. The Wnt effector β-catenin is required for hair cell specification and has been shown to directly activate *Atoh1* transcription in neuronal progenitors (Shi et al., 2010; Shi et al., 2014). Our here presented data is most consistent with a model in which Wnt/β-catenin signaling is required to initiate *Atoh1* expression, whereas Activin signaling functions to enhance *Atoh1* expression/activity, providing spatial and temporal control over hair cell differentiation. Wnt/β-catenin signaling and TGF-β-related pathways are known to cooperate with the transcriptional regulation of their target genes (Labbe et al., 2000) (Luo, 2017) and it will be of interest to resolve whether and how Activin and Wnt signaling pathways collaborate in *Atoh1* gene regulation. Furthermore, it will be of interest to identify the upstream signals/factors that induce *Inhba* expression at the onset of hair cell differentiation. A potential candidate is the Notch signaling pathway. Notch signaling is active in pro-sensory cells and has been recently shown to positively regulate *Inhba* expression in differentiating and terminal differentiated supporting cells (Campbell et al., 2016; Maass et al., 2016).

### FST maintains pro-sensory cells in an undifferentiated and proliferative state

We show that FST overexpression severely delays the onset of pro-sensory cell cycle withdrawal and differentiation. A similar function in timing pro-sensory cell withdrawal and differentiation has been recently reported for the RNA binding protein LIN28B (Golden et al., 2015). Interestingly, the expression of *Fst* in the developing cochlea largely mimics that of *Lin28b;* each are initially highly expressed in pro-sensory cells and are downregulated upon differentiation following a basal-to-apical gradient. Regulatory and functional connections between Activin signaling and LIN28B have not yet been established and it will be of interest to determine whether a link between LIN28B and FST exists. TGFβ-related signaling pathways are known to inhibit proliferation by increasing the expression and/or activity of cyclin dependent kinase (CDK) inhibitors (Massague and Gomis, 2006). In the developing cochlea, the CDK inhibitor p27/Kip1 (CDKN1B) is the main regulator of pro-sensory cell cycle exit and its transcriptional upregulation directs the apical-to-basal wave of pro-sensory cell cycle withdrawal (Chen and Segil, 1999; Lee et al., 2006). However, we found no evidence that FST overexpression interfered with the transcriptional regulation of *p27/Kip1*, suggesting that the observed delay in pro-sensory cell cycle exit was likely caused by a reduction in P27/Kip1 protein stability and/or activity. In addition, it is likely that the increase in *Id3* and *Id4* expression in response to FST overexpression contributed to the observed delay in pro-sensory cell cycle exit. High ID protein expression is associated with a proliferative, undifferentiated cell state and ID4 expression in spermatogonia and mammary glands is a predictor for stemness (Wang and Baker, 2015; Helsel et al., 2017).

### A radial Activin-FST counter gradient controls the production of inner hair cells

Interestingly, we find that FST overexpression and to a lesser extend ablation of Activin A (*Inhba*) results in an overproduction of inner hair cells. Based on our findings we propose that FST overexpression disrupts a previously unrecognized radial gradient of pro-sensory cell cycle withdrawal, leading to prolonged inner hair cell progenitor proliferation. We provide evidence for the existence of a radial (medial-to-lateral) gradient of Activin A activity. We show that at the peak of terminal mitosis (E13.5), as *Inhba* expression is induced within the basal pro-sensory domain, *Fst* expression is maintained at the lateral edge of the pro-sensory domain, creating a medial-to-lateral gradient of Activin A signaling. Furthermore, we show that in wild type tissue inner hair cell progenitors withdraw from the cell cycle prior to outer hair cell progenitor cells and demonstrate that this distinct medial-to-lateral gradient of pro-sensory cell cycle withdrawal is disrupted in response to FST overexpression. This newly identified radial gradient of terminal mitosis constitutes a novel mechanism for limiting the number of inner hair cells being produced in the mammalian cochlea.

## METHODS

### Experimental animals

Atoh1/nEGFP transgenic mice (Lumpkin et al., 2003) were obtained from Jane Johnson, University of Texas, Southwestern Medical Center. Pax2-Cre BAC transgenic mice (Ohyama and Groves, 2004) were obtained from Andrew Groves, Baylor College. Inhba floxed mice (Pangas et al., 2007) were obtained from Martin Matzuk, Baylor College. The R26-M2rtTA (Hochedlinger et al., 2005)(stock no. 006965) was purchased from Jackson Laboratories (Bar Harbor, ME). FST transgenic mice were obtained from Se-Jin Lee, Johns Hopkins University, School of Medicine. In this line a cassette encoding the human FST-288 isoform is under the control of a tetracycline-responsive promoter element (tetO) (Lee, 2007; Roby et al., 2012). Mice were genotyped by PCR. Pax2-Cre: Cre1F (GCCTGCATTACCGGTCGATGCAACGA), Cre1R (GTGGCAGATGGCGCGGCAACACCATT) yields a 700bp band. Inhba floxed: Inhba fx1 (AAG AGA GAA TGG TGT ACC TTC ATT), Inhba fx2 (TAT AAC CTG GGT AAG TGG GT), Inhba fx3 (AGA CGT GCT ACT TCC ATT TG) yield a 400bp band for the floxed allele and a 280bp for the wild type allele. R26-M2rtTA: MTR (GCG AAG AGT TTG TCC TCA ACC), F (AAA GTC GCT CTG AGT TGT TAT), WTR (GGA GCG GGA GAA ATG GAT ATG) yield a 340bp band for the mutant allele and a 650bp for the wild type allele. FST: YA88 (TTGCCTCCTGCTGCTGCTGC), YA123 (TTTTTCCCAGGTCCACAGTCCACG) yields a 247bp band for the FST transgene.

Atoh1/nGFP: EGFP1 (CGA AGG CTA CGT CCA GGA GCG CAC), EGFP2 (GCA CGG GGC CGT CGC CGA TGG GGG TGT) yields a 300bp band for EGFP. Mice were maintained on a C57BL/6; CD-1 mixed background. Mice of both sexes were used in this study. Embryonic development was considered as E0.5 on the day a mating plug was observed. To induce *FST* transgene expression doxycycline (dox) was delivered to time-mated females via ad libitum access to feed containing 2 grams of dox per kilogram feed (Bioserv no. F2893). All experiments and procedures were approved by the Johns Hopkins University Institutional Animal Care and Use Committee (protocol #MO17M318), and all experiments and procedures adhered to the National Institutes of Health-approved standards.

### Tissue harvest and processing

Embryos and early postnatal pups were staged using the EMAP eMouse Atlas Project (http://www.emouseatalas.org) Theiler staging criteria. Inner ear cochleae were collected in Hanks buffer (Corning Cellgro). To free the cochlear epithelial duct from surrounding tissue, dispase (1mg/ml; Invitrogen) and collagenase (1mg/ml; Worthington) mediated digest was used as previously described (Golden et al., 2015). To obtain cochlear whole mount preparations (also referred to as surface preparations) containing the auditory sensory epithelium (E18.5-P0), the cochlear capsule, spiral ganglion, and Reissner’s membrane were removed, and the remaining tissue was briefly fixed in 4% (vol/vol) paraformaldehyde (PFA) (Electron Microscopy Sciences) in PBS. To obtain cochlear sections, heads were fixed in 4% PFA in PBS, cryoprotected using 30% sucrose in PBS, and embedded in OCT (Sakura Finetek). Tissue was sectioned at a thickness of 14 μm and collected on SuperFrost Plus slides (Thermo Scientific) and stored at −80°C.

### Histochemistry and in situ hybridization

Immunostaining was performed according to the manufacturer’s specifications. Primary antibodies: rabbit anti-myosinVIIa (1:500, Proteus no. 25-6790), goat anti-SOX2 (1:500, Santa Cruz no. sc-17320). Cell nuclei were fluorescently labeled with Hoechst-33258 dye (Sigma). Actin filaments were labeled with Alexa Fluor (488 or 546) conjugated phalloidin (1:1000, Invitrogen). Alexa Fluor (488 or 546) labeled secondary antibodies (1:1000, Invitrogen) were used. For insitu hybridization digoxigenin (DIG)-labeled antisense RNA probes were prepared according to the manufacturer’s specifications (Roche). PCR amplified fragments of *Inhba* (NM_008380, 47-472) and *Fst* (NM_008046, 180-492) were used as template and gene-specific T7 RNA polymerase promoter hybrid primers were used for in vitro transcription. *Atoh1* (NM_007500) and *Sox2* (NM_011443) probes were prepared as previously described (Golden et al., 2015). Probes were detected with the anti–DIG-AP (alkaline phosphatase) conjugated antibody (Roche), and the color reactions were developed by using BM Purple AP Substrate (Roche).

### Cochlear explant culture and hair cell differentiation assay

Wild type or FST overexpressing embryos and their control littermates were screened for native EGFP (Atoh1/nEGFP) expression and staged (see tissue harvest and processing). Embryos of inappropriate stage and Atoh1/nEGFP negative embryos were discarded. Cochleae from individual embryos were harvested in Hanks media (Life Technologies), and treated with dispase and collagenase to remove cochlear capsule. The remaining tissue, including the cochlear epithelial duct, the vestibular sacculus, and the innervating spiral ganglion, was placed onto filter membranes (SPI Supplies, Structure Probe) and cultured in DMEM-F12 (Life Technologies), 1% FBS (Atlanta Biologicals), 5 ng/ml EGF (Sigma), 100 U/ml penicillin-streptomycin (Sigma), and 1xB27 supplement (Life Technologies). Activin A (final conc. 500ng/ml) was added at plating and was replenished daily. All cultures were maintained in a 5% CO_2_ / 20% O_2_ humidified incubator. To monitor hair cell differentiation, green fluorescent images of native EGFP expression were captured using fluorescent stereo-microscopy (Leica). Fluorescent images were analyzed in Photoshop CS6 (Adobe), and lengths of Atoh1/nEGFP-positive domains were measured using ImageJ software (National Institutes of Health).

### Recombinant protein

Recombinant human/mouse/rat Activin A (R&D Systems no. 338-AC) was reconstituted in sterile PBS containing 0.1% BSA at a concentration of 50 μg/ml and used at final concentration of 200-500 ng/ml. Recombinant human BMP4 (R&D Systems, no. 314-BP-010) was reconstituted in sterile 4 mM HCl 0.1% BSA at a concentration of 100 μg/ml and used at 100 ng/ml final concentration. Stock solutions were stored at −80°C.

### RNA extraction and q-PCR

Cochlear epithelia were isolated from cultured cochlear explants or freshly harvested inner ear tissue using dispase/collagenase treatment. RNeasy Micro kit (Qiagen) was used to isolate total RNA, and mRNA was transcribed into cDNA using iScript kit (Bio-Rad). Q-PCR was performed with Fast SYBR Green Master Mix reagent (Applied Biosystems) and gene-specific primer sets on a CFX-Connect Real Time PCR Detection System (Bio-Rad). Each PCR was performed in triplicate. Relative gene expression was analyzed by using the ΔΔCT method (Schmittgen and Livak, 2008). The ribosomal gene Rpl19 was used as endogenous reference gene. The following q-PCR primers were used:

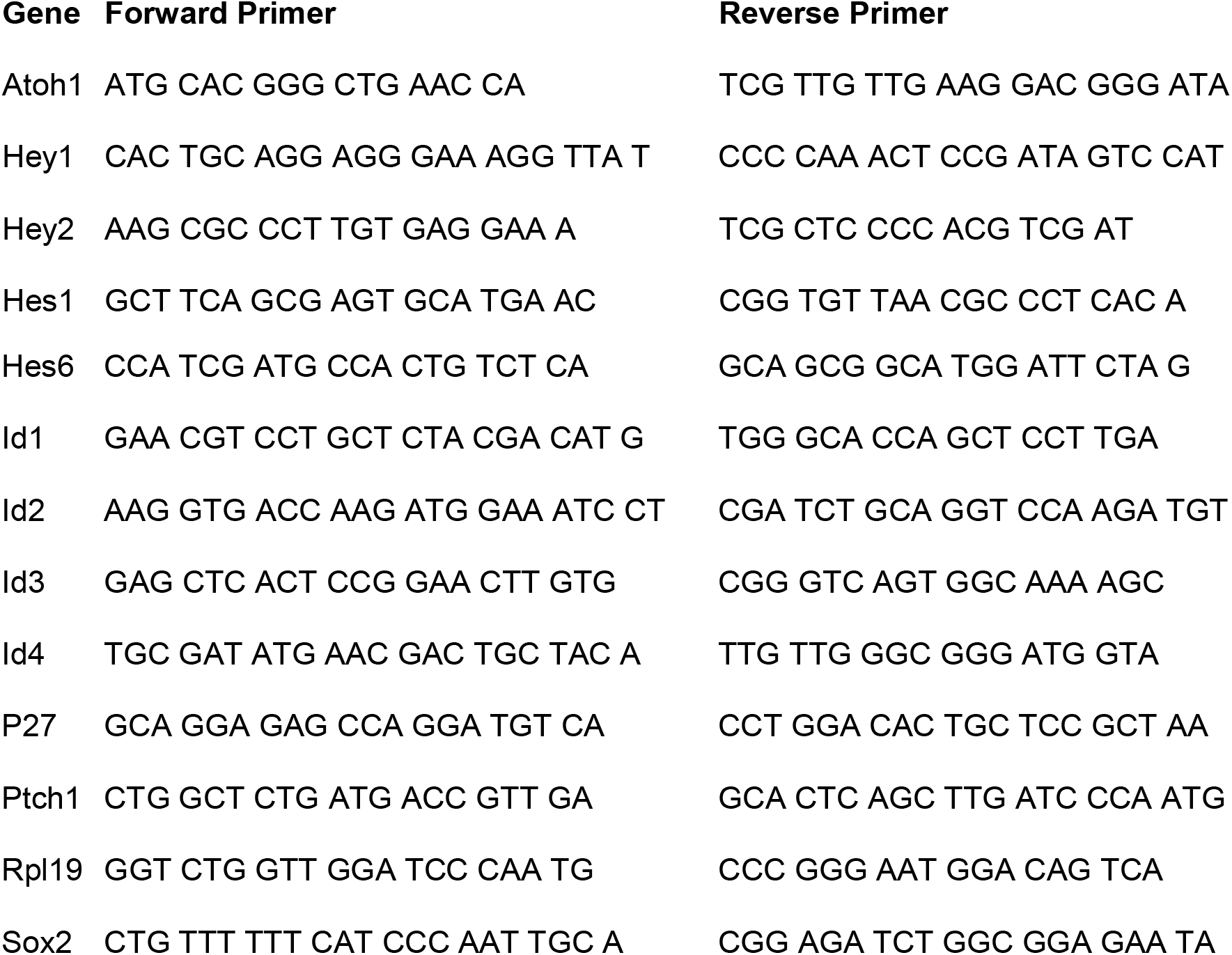

### Western blot

Individual cochlear epithelia were lyzed in RIPA lysis buffer (Sigma) supplemented with Roche Protease Inhibitor (Sigma) and Phosphatase Inhibitor Cocktail no.2 and no.3 (Sigma). Following manufactures recommendations, equal amounts of cochlear protein extract were resolved on NuPAGE 4-12% Bis-Tris Gels (Invitrogen) and transferred to Immun-Blot PVDF membrane (Bio-rad) by electrophoresis. Membranes were blocked in 5% nofat dry milk in TBST and immunoblotted with rabbit anti-P-Smad2/3 (1:1,000 Cell Signaling, no.8828), P-Smad1/5/9 1:1,000 (Cell Signaling, no.13820) and mouse anti-β-actin 1:1,000 (Santa Cruz, no.SC-47778). HRP-conjugated secondary antibodies from Jackson Immuno Research were used at a concentration of 1:10,000 (goat anti-rabbit IgG, no.111-035-003; sheep anti-mouse IgG no.515-035-003). Signal was revealed using a Western Lightening-ECL kit (Perkin Elmer) or SuperSignal West Femto Maximum Sensitivity Substrate (Thermo Scientific) according to manufacturer’s instructions.

### Quantification of hair cells and supporting cells

Cell counts were performed in cochlear whole mounts. Hair cells were identified by their native EGFP (Atoh1/nEGFP) expression, as well as by immuno-staining for the hair cell-specific protein myosinVIIa (Myo7a); supporting cells were identified by SOX2 immuno-staining. Classification of cell subtypes was based on their location within the sensory epithelium. Low-power confocal and epi-fluorescent images (Zeiss) of the hair cell layer were used to reconstruct the entire cochlear sensory epithelium. The resulting composite images were used to count ectopic inner hair cells, measure the total length of the sensory epithelia and used to define basal, mid and apical segments (~1400 μm). The apical tip (~300 – 500 μm) was excluded from the analysis. For hair cell and supporting cell counts a series of high-power confocal (Zeiss) z-stack images spanning the hair cell and supporting cell layer were taken within the basal, mid and apical segments. Images were assembled and analyzed in Photoshop CS6 (Adobe). Image J software (National Institutes of Health) was used to measure the length of counted segments and total length of the sensory epithelium.

### Proliferation assay

EdU (5-ehynyl-2’-deoxyuridine, Invitrogen) was reconstituted in PBS and administered at 50 μg per gram of body weight to time-mated pregnant dams per intraperitoneal injection. Click-iT AlexaFluor-488 or −546 Kit (Invitrogen) was used to detect incorporated EdU according to the manufacturer’s specifications. To quantify the number and percentage of EdU positive hair cells and supporting cells, EdU stained cochlear surface preparations were co-stained with the nuclear dye Hoechst-33258 (Sigma) and immuno-stained for SOX2, and Myo7a. Length measurement and cell counts were conducted as described above.

### Statistical reporting

Values are presented as mean ± standard error of the mean (SEM). The sample size (n) represents the number of biological independent samples (biological replicates) analyzed per experimental group. Two-tailed unpaired Student’s tests were used to determine the confidence interval p-values ≤ 0.05 were considered significant. P-values > 0.05 were considered not significant. Biological independent samples (biological replicates) were allocated into experimental groups based on genotype and/or type of treatment. A minimum of three biological independent samples were analyzed per group. To avoid bias masking was used during data analysis.

## ACKNOWLEDGMENTS

We thank the members of the A.D. Laboratory and the Center for Sensory Biology for the help and advice provided throughout the course of this study. We thank Jane Johnson for the Atoh1/nEGFP mice; Andrew Groves for the *Pax2-Cre* mice; Se-Jin Lee for *FST* transgenic mice and Martin Matzuk for *Inhba* floxed mice. This work was supported by National Institutes of Health Grants DC013477 (A.B.-G.) DC012972 (E.J.G.), DC016538 (M.P.), DC011571 (A.D.), DC005211 (Sensory Mechanisms Research Core Center) and David M. Rubenstein Fund for Hearing Research (A.D.).

